# Age and Visual Contrast Affect Walking Stability on Winding Paths

**DOI:** 10.64898/2026.11.17.767409

**Authors:** Jonathan B. Dingwell, Carlie J. Hernjak, Sarah E. Overby, Tarkeshwar Singh

**Affiliations:** Department of Kinesiology, The Pennsylvania State University, University Park, PA 16802 USA

## Abstract

Older adults often experience loss of balance, and potentially falls, when trying to maintain lateral stability while walking. Age-related declines in vision likely exacerbate these challenges. Goal-directed walking to navigate complex environments requires that we clearly see the task-relevant features of our environment, known as visual salience. This study quantified lateral stability of 16 young (10F/6M; Age 22.8±1.95 yrs) and 16 older (12F/4M; Age 70.3±1.82 yrs) healthy adults walking on a treadmill within a Motek M-Gait virtual reality system. Participants walked on either straight or continuously-winding paths, each presented with either high or low visual contrast (salience). We quantified mean lateral margins of stability (*μ(MoS*_*L*_)), *MoS*_*L*_ variability (*σ*(*MoS*_*L*_)), and lateral probability of instability (*PoI*_*L*_). We also quantified lateral base of support width (*BoS*_*w*_) and percent of unstable steps. Straight walking elicited few differences between groups or visual contrast conditions. Conversely, participants walking on low contrast winding paths took narrower steps and exhibited correspondingly decreased *μ(MoS*_*L*_), but also exhibited increased *σ*(*MoS*_*L*_) and greatly increased *PoI*_*L*_. Thus, despite the high and low contrast winding paths being biomechanically equivalent, people chose to take narrower steps on the harder-to-see low contrast paths, despite such narrower steps increasing their risk of lateral instability. Conversely, we observed few group (young vs. older) differences, and no significant group by contrast interaction effects. Thus, these older adults may have adopted effective compensatory stepping adjustments to maintain their stability. Our findings suggest that fall prevention interventions might benefit from manipulating visual uncertainty about stepping/walking task goals.

## INTRODUCTION

Maintaining side-to-side balance during everyday walking is essential for safe mobility, particularly as individuals navigate complex environments. As physical and cognitive function decline as we age, our ability to adapt stepping and maintain stability becomes compromised, contributing to the high prevalence of falls among older adults (Fischer et al., 2014; Kakara et al., 2023). Indeed, non-fatal falls among older adults incurred $80 billion in healthcare expenditures in 2020 (Haddad et al., 2024). Notably, improper weight shifting during forward walking is a common cause of falls (Robinovitch et al., 2013), emphasizing the need for more effective assessments of lateral dynamic balance and fall risk in older adults (Cofré Lizama et al., 2015; Yang et al., 2020).

During forward walking, humans are inherently more unstable in the mediolateral direction (Bauby and Kuo, 2000; Kuo, 1999). Hence, lateral balance is thought to require active control via sensory feedback (Bruijn and van Dieën, 2018). One widely used measure of lateral stability is the lateral margin of stability (*MoS*_*L*_) (Hof, 2008; Hof et al., 2005). While most studies average *MoS*_*L*_ over multiple steps or strides (Watson et al., 2021), computing such averages obscures important step-to-step variability in balance strategies, especially as stability must be re-established with each new step (Kazanski et al., 2022). To address this, we previously introduced the lateral Probability of Instability (*PoI*_*L*_) that directly quantifies the likelihood that *MoS*_*L*_ will be negative (i.e., unstable) for any given step across a range of steps (Kazanski et al., 2022).

Additionally, walking in the real world involves a substantial amount of *non*-straight walking (Nikmanesh et al., 2025; Wang and Adamczyk, 2019), as we navigate cluttered environments and avoid obstacles in our path, be they fixed (e.g., furniture) or moving (i.e., other people). Older adults are less able to shift weight effectively (Ambrose et al., 2013). This impairs their ability to enact appropriate stepping maneuvers in walking tasks that require such weight shifting (Desmet et al., 2024; Yamada et al., 2012). They also make riskier collision-avoidance maneuvers when encountering other moving people (Rapos et al., 2021). These impaired capacities may contribute to their increased fall risk (Robinovitch et al., 2013). When young adults walk on continuously-winding paths, they take narrower steps and make more lateral stepping adjustments (Render et al., 2025). Conversely, on slowly-narrowing paths, older adults take wider steps, which may help preserve stability (Kazanski et al., 2024).

Vision deteriorates as we age (Owsley, 2011) and this impaired vision contributes to falls (Black and Wood, 2005; Saftari and Kwon, 2018). Vision is crucial for walking (Tong et al., 2017), especially for guiding how we navigate the real world (Muller et al., 2024), and these processes require integration across multiple cortical areas (Drew and Marigold, 2015).

Our brains use visual information to make anticipatory gait adjustments 1-3 steps in advance (Marigold and Patla, 2007; Matthis et al., 2018; Patla and Greig, 2006). Age-related declines in visual processing and saliency detection impair performance during visual-search tasks (Madden, 2007). While walking, older adults visually sample their environments differently (Dowiasch et al., 2015) in ways that *may* impact their balance (Meissner et al., 2025). Yet, few studies have systematically quantified how visual conditions influence lateral balance during complex walking tasks. In particular, how the visual saliency of one’s walking path affects one’s lateral stability has not been characterized in older adults.

Therefore, this study determined how the visual saliency of one’s walking path affects lateral stability in both young and older adults walking on both straight and winding paths. We quantified the Probability of Instability (*PoI*_*L*_) to assess step-to-step risk of balance loss, providing a sensitive metric of lateral balance control. We hypothesized that as visual information was diminished, *PoI*_*L*_ would increase for all participants. We further hypothesized that older adults would be disproportionately affected by decreased walking path visual salience. By further elucidating how visual and mechanical demands interact to influence balance, this work contributes to a more comprehensive understanding of age-related fall risk and offers insight into designing better prevention strategies for at-risk populations.

## METHODS

### Participants

We recruited 25 young (YH; age 18-35) and 31 older (OH; age ≥ 65) healthy adults. All potential participants provided written informed consent as approved by the Institutional Review Board at The Pennsylvania State University. We excluded individuals if they had blood pressure >140/90 mmHg (Frese et al., 2011), body mass index (BMI) <18 or >30, or corrected visual acuity worse than 20/40 (Fabre et al., 2010).

Potential participants then completed a 5-Meter Walk Test (5mWT) (Ng et al., 2013), Timed Up and Go (TUG) test (Bohannon, 2006), and Four Square Step Test (FSST) (Dite and Temple, 2002; Moore and Barker, 2017). We excluded individuals if they had walking speed <0.8m/s from the 5mWT, TUG times >12 sec, or FSST times >15 sec.

We assessed perceived balance confidence using the Iconographic Falls Efficacy Scale (I-FES) (Delbaere et al., 2011; Lim et al., 2021). We used the Freiburg Visual Acuity and Contrast Test (FrACT) (Bach, 1996; Render et al., 2026) to ensure all participants had visual contrast sensitivity within normal ranges (Bartholomew et al., 2016; Friedel et al., 2025).

Twenty participants in each group passed screening and completed the full experiment. Due to technical issues with data collection, final data analyses were performed for 16 participants in each group (Fig. 1).

**Figure 1.**
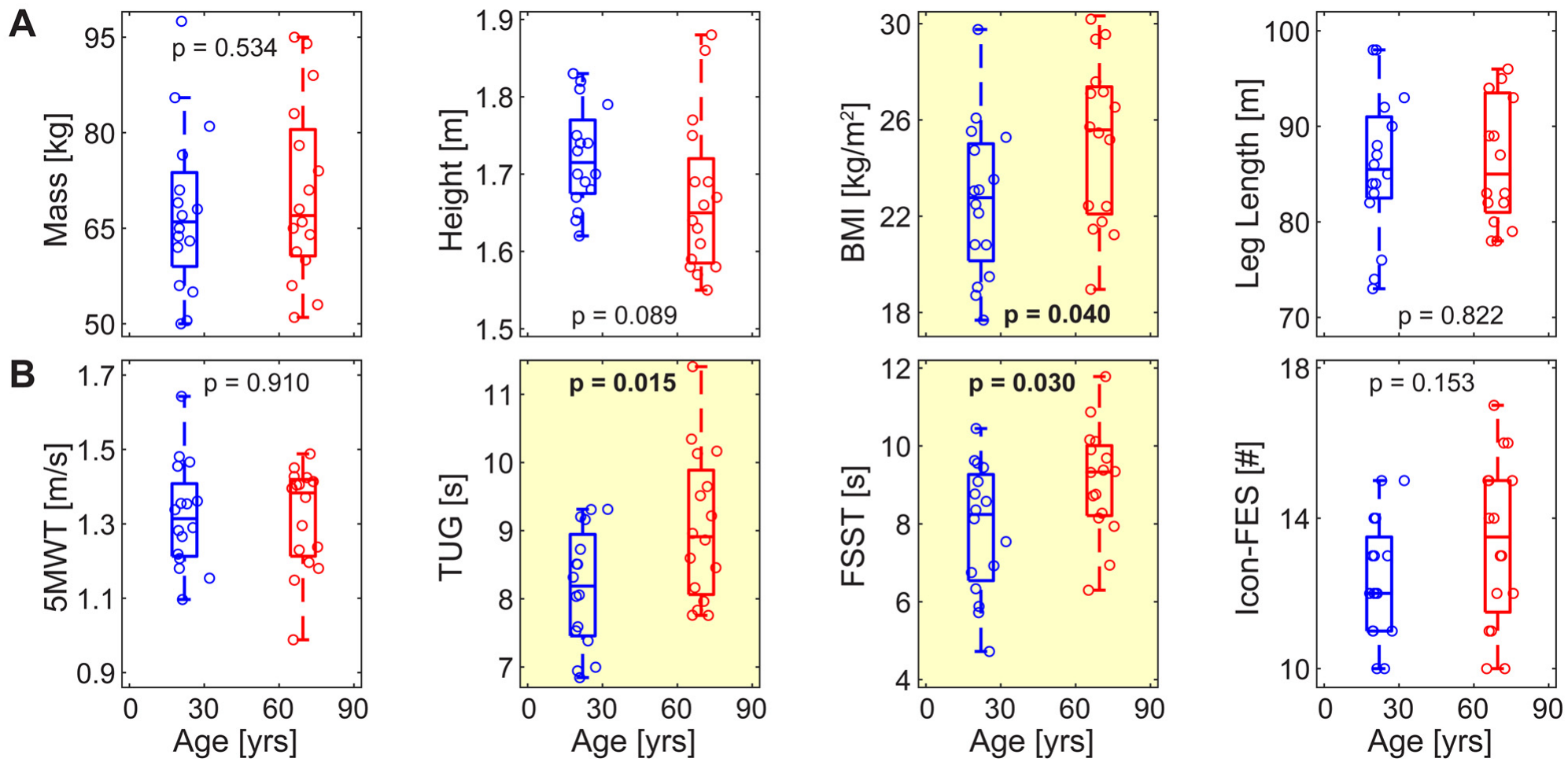
Participant Characteristics and Assessment Scores. (**A**) Demographics (body mass, height, BMI, and leg length) for each participant and group, plotted versus their age. Individual markers represent individual participants in each group (YH blue; OH red). Box plots are plotted at the mean age for each group. Each box plot shows the median, 1^st^ and 3^rd^ quartiles, and whiskers extending to 1.5× the inter-quartile range for that group. (**B**) Baseline assessments (5MWT, TUG, FSST, and Icon-FES) for each participant and group, plotted in the same manner as in (**A**). P-values are shown in each subplot and significant (p < 0.05) Group differences are highlighted in yellow.

### Protocol and Data Collection

Participants walked on a 1.2 m-wide treadmill within a 240° curved screen VR system (Motek M-Gait, Amsterdam, Netherlands) (Fig. 2A). They wore a safety harness and appropriate walking shoes. A forest scene was projected. Stepping off the virtual path triggered visual (fireworks) and auditory (flute) feedback. Each participant wore 18 retroreflective markers, including 5 on their head, 1 on their right shoulder, 4 around their pelvis (left & right ASIS & PSIS), and 4 on each foot (1^st^ & 5^th^ metatarsal heads, lateral malleolus, and calcaneus).

**Figure 2.**
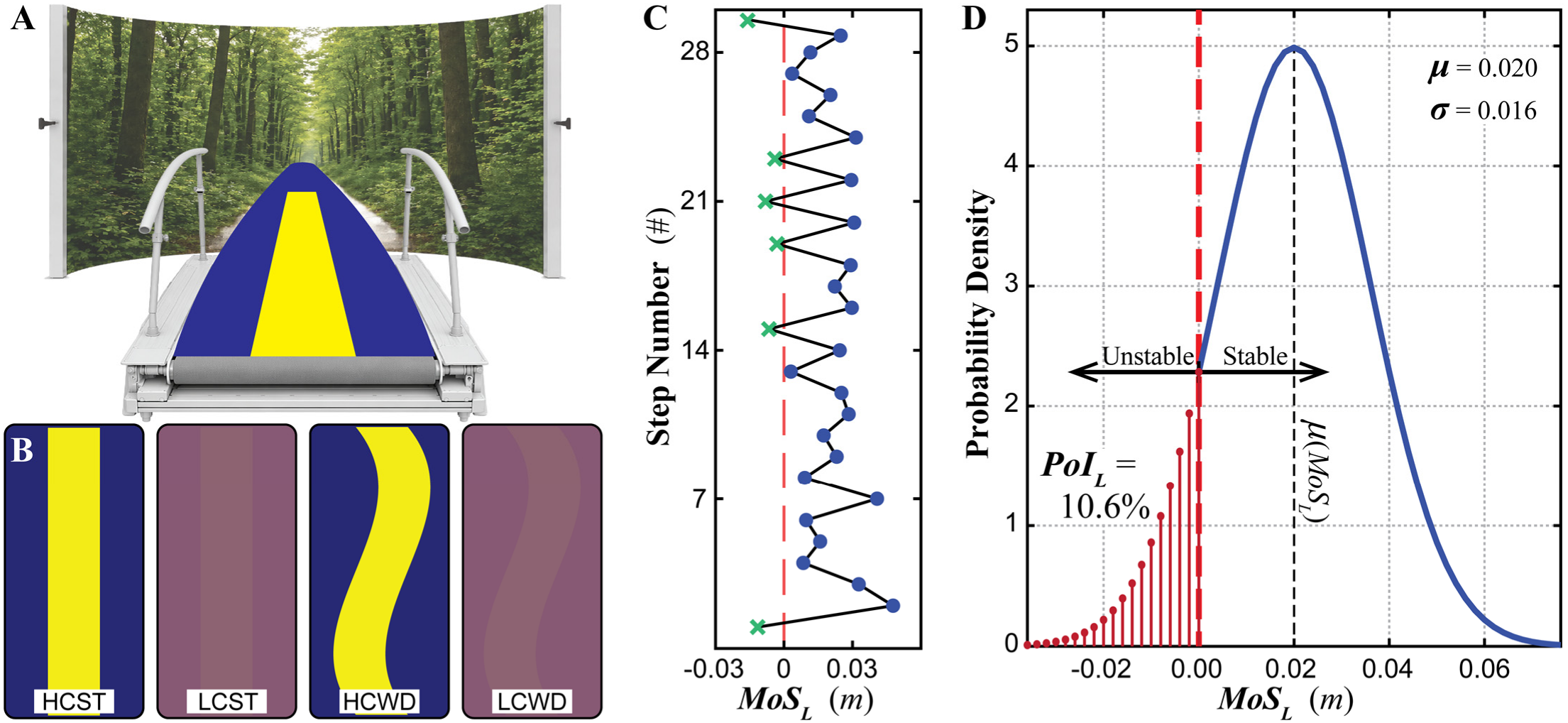
Experiment and Primary Analyses. (**A**) Schematic of experimental setup. (**B**) Schematic representation of the conditions tested: two Straight (ST) walking paths of either High (HC) or Low (LC) contrast, and two Winding (WD) walking paths of either HC or LC. (**C**) Schematic representation of step-to-step margin of stability (*MoS*_*L*_) data, where steps categorized as “stable” (i.e., *MoS*_*L*_ > 0) are shown as blue dots (●) and those categorized as “unstable” (i.e., *MoS*_*L*_ < 0) are shown as green crosses (×). (**D**) Schematic representation of how probability of instability (*PoI*_*L*_; Eq. 3) is calculated from a hypothetical full distribution of such *MoS*_*L*_ values (Kazanski et al., 2022). *PoI*_*L*_ (here, 10.6%) directly calculates the cumulative probability (area under the curve) of taking an unstable (*MoS*_*L*_ < 0) step (here, 1 unstable step every ~ 9.4 steps). Conversely, the mean (here, *μ*(*MoS*_*L*_) = +0.020 m) yields *no* information regarding those unstable steps. Hence, *PoI*_*L*_ directly estimates statistical risk of instability, consistent with Hof’s theory (Hof et al., 2005), whereas *μ*(*MoS*_*L*_) does not.

Participants walked on 0.45 m-wide projected paths that varied in lateral oscillation and color contrast (color difference between path and floor). Paths were generated as (Render et al., 2024; Render et al., 2027):

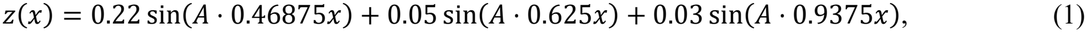

where *z* (m) is the lateral position of the path center, *x* (m) is forward distance, and *A* is a frequency scaling factor. We set *A*=0 for Straight (ST) paths, *A*=1 for slowly-winding paths (used for acclimation only), and *A*=3 for Winding (WD) paths.

High Contrast (HC) paths projected a bright yellow (RGB = [254, 255, 0]) path onto a dark blue (RGB = [0, 0, 129]) background (Fig. 2A). Medium contrast paths (used for acclimation only) projected a purple (RGB = [152, 113, 109]) path onto a darker (RGB = [115, 66, 120]) background. Low Contrast (LC) paths projected a somewhat darker purple (RGB = [144, 102, 112]) path onto a somewhat lighter purple (RGB = [132, 86, 116]) background (Fig. 2B) (Render et al., 2026).

To avoid introducing confounding effects of different speeds (Ettema et al., 2025; Fukuchi et al., 2019), all walking were conducted at 1.0 m/s, consistent with preferred treadmill (McDonald et al., 2026) and overground (Brinkerhoff et al., 2022) walking speeds. Holding speed constant also ensured an inertial reference frame moving with the treadmill belt, so walking dynamics would be comparable to walking overground (Jung and Koo, 2022; van Ingen Schenau, 1980).

Each participant first completed an acclamation protocol involving six 1½-minute walking trials that progressed in difficulty: from full-width (1.2m wide) path, to Straight (0.45m wide) HC path (HCST; Fig. 2B), to slowly-winding HC path, to full Winding HC path (HCWD; e.g., Fig. 2A-B), to medium contrast Winding path, and finally to low contrast Winding path (LCWD; Fig. 2B).

We then asked each participant to complete one 1-minute acclimation trial and two 4-minute test trials for each of four test conditions: (Straight, Winding) × (HC, LC). We randomized the order-of-presentation of these conditions for each participant, while counterbalancing presentation order across participants using a Latin square design. Participants rested between trials and were instructed to maintain balance and remain on the path at all times. For each 4-minute experimental trial, we collected kinematic data at 100 Hz using a 9-camera Vicon motion capture system (Oxford Metrics, UK) integrated into the M-Gait VR system.

Importantly, because each path (Straight or Winding) was the same, and because we held speed constant, the mechanical requirements to walk on each path were exactly the same for both path Contrasts (Render et al., 2027). Thus, there was biomechanical reason for participants not to be able to walk with the same stepping mechanics and balance control on both HC and LC paths.

### Data Processing and Analyses

We pre-processed raw marker data in Vicon Nexus. We then exported final marker trajectory data and path motion data from D-flow to analyze in MATLAB (MathWorks, Natick, MA).

We low-pass filtered marker trajectories with a zero-lag 4^th^-order Butterworth filter (cutoff 10 Hz) and interpolated these data to 600 Hz (Bohnsack-McLagan et al., 2016) to improve precision in identifying stepping events. We identified individual steps using a validated heel-pelvis distance algorithm (Zeni et al., 2008). We then transformed marker location data into a local path-aligned coordinate system (Dingwell et al., 2023) to account for direction changes along the winding path (Render et al., 2024). We analyzed the first 340 consecutive steps of each trial.

We quantified lateral stability at each step from the lateral Margin of Stability, *MoS*_*L*_ (Hof, 2008; Hof et al., 2005; Kazanski et al., 2022). For each step, we took the locally-lateral (*z*) coordinate of the leading foot’s 5^th^ metatarsal marker as the lateral edge of the base of support (*u*_*max*_). We used pelvic centroid (average of 4 pelvis markers) motion to define the locally-lateral center-of-mass (CoM) state (*z, ż*) (Havens et al., 2018). We calculated the *minimum MoS*_*L*_ for each step as:

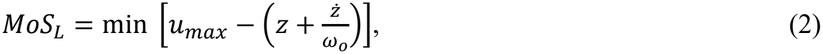

where 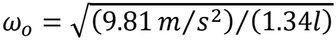, and *l* was leg length (Kazanski et al., 2022). Here, a step is considered “stable” if *MoS*_*L*_ > 0, or “unstable” if *MoS*_*L*_ < 0 (Fig. 2C). For each trial of 340 steps, we computed the mean, *μ*(*MoS*_*L*_), and standard deviation, σ(*MoS*_*L*_). From these, we then computed the lateral Probability of Instability (*PoI*_*L*_) (Kazanski et al., 2022):

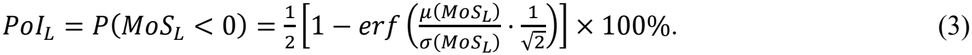

Whereas *μ*(*MoS*_*L*_) and σ(*MoS*_*L*_) calculate the mean and variance of *all* steps (stable and unstable), *PoI*_*L*_ directly calculates the cumulative probability of *only* unstable (*MoS*_*L*_ < 0) steps occurring (Fig. 2D). Thus, *PoI*_*L*_ is consistent with Hof’s original theory of instability, whereas *μ*(*MoS*_*L*_) is not (Kazanski et al., 2022).

In addition, we also calculated the base-of-support width, *BoS*_*w*_, for each step as the locally-lateral (*z*-direction) absolute distance between the 5^th^ metatarsal markers of each foot (Kazanski et al., 2024). We then computed the mean, *μ*(*BoS*_*w*_), across all steps for each trial. Lastly, to compare with *PoI*_*L*_ (Render et al., 2024), we directly computed the percent of unstable (i.e., *MoS*_*L*_ < 0; e.g., Fig. 2C) steps in each trial.

### Statistical Analyses

We analyzed data for Straight and Winding paths separately (Render et al., 2024, 2025). The dependent measures included *BoS*_*w*_, % Unstable Steps, *µ*(*MoS*_*L*_), *σ*(*MoS*_*L*_), and *PoI*_*L*_. For each, we applied a mixed-effects analysis of variance (ANOVA) with repeated measures. Fixed effects were Group × Contrast. Subject (nested within Group) was included as an additional random factor. When indicated, we first log-transformed some dependent measures to satisfy normality and linearity assumptions of the ANOVA. When we found significant Group × Contrast interactions, we performed Tukey’s pairwise post-hoc comparisons to test for differences between Groups (YH vs. OH) for each path Contrast (HC vs. LC), and for differences between path Contrasts for each Group.

## RESULTS

### Straight Paths

When walking on Straight paths, participants took mostly stable (*MoS*_*L*_ > 0) steps, with few if any unstable (*MoS*_*L*_ < 0) steps (e.g., Fig. 3A), as expected (Render et al., 2024; Render et al., 2027).

**Figure 3.**
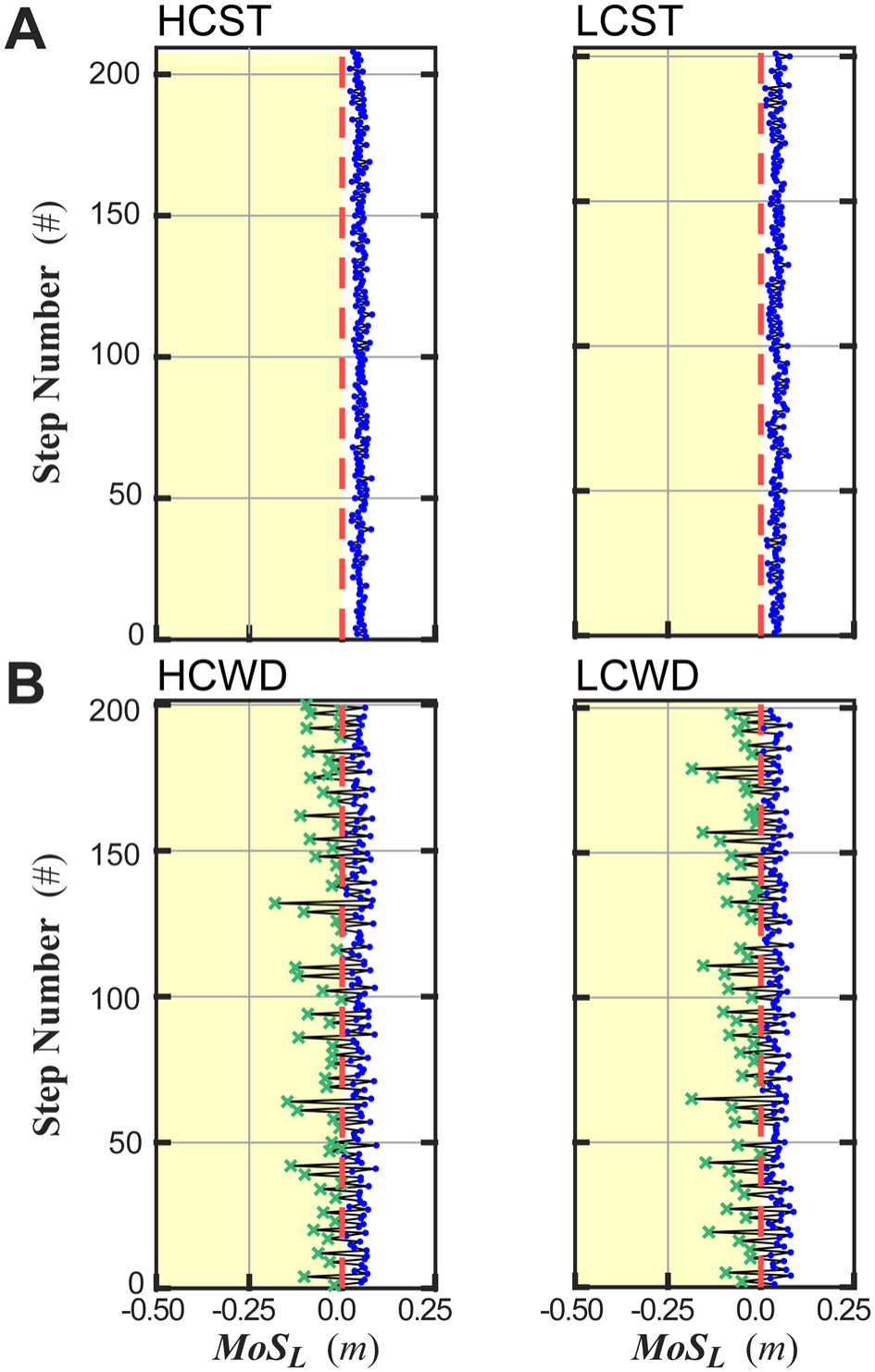
Example *MoS*_*L*_ Time Series Data. Example lateral Margin of Stability (*MoS*_*L*_) values at each consecutive step for ~200 steps each from each of four representative trials from a typical (OH) participant walking (**A**) on Straight paths of both high (HC) and low (LC) visual contrast, and (**B**) on Winding paths of both high (HC) and low (LC) visual contrast. In each subplot, the vertical red dashed line indicates *MoS*_*L*_ = 0. The shaded yellow region highlights the range where steps would be considered “unstable” (i.e., *MoS*_*L*_ < 0). Steps where *MoS*_*L*_ > 0 are shown as blue dots (●), while steps where *MoS*_*L*_ < 0 are shown as green crosses (×).

While walking on Straight paths, both YH and OH participants exhibited similar *μ*(*MoS*_*L*_) on paths of both contrasts (Fig. 4A; all p ≥ 0.089; Table 1). For *σ*(*MoS*_*L*_), participants in both groups exhibited greater step-to-step variability on low (LC) compared to high (HC) paths (Fig. 4B; p = 0.008; Table 1). However, there were no significant differences between either Groups or Contrasts for *PoI*_*L*_ (Fig. 4C; all p ≥ 0.229; Table 1). There were no significant Group × Contrast interactions for any of these measures (all p ≥ 0.438; Table 1).

**Table 1.** Statistical Results for Walking on Straight Paths. Differences between age Group (YH vs. OH) and path visual Contrast (HC vs. LC) for the data shown in Figs. 4 and 5.

| Data In | Variable | Group | Contrast | Group ×<br>Contrast |
|---|---|---|---|---|
| Fig. 4 [A]: | *μ*(*MoS<sub>L</sub>*) | F<sub>(1,30)</sub> = 3.01 | F<sub>(1,94)</sub> = 2.96 | F<sub>(1,94)</sub> = 0.36 |
|  |  | p = 0.093 | p = 0.089 | p = 0.549 |
| [B]: | ln(*σ*(*MoS<sub>L</sub>*)) | F<sub>(1,30)</sub> = 1.30 | F<sub>(1,94)</sub> = 7.33 | F<sub>(1,94)</sub> = 0.61 |
|  |  | p = 0.263 | **p = 0.008** | p = 0.438 |
| [C]: | ln(*PoI<sub>L</sub>*) | F<sub>(1,30)</sub> = 0.07 | F<sub>(1,94)</sub> = 1.46 | F<sub>(1,94)</sub> = 0.40 |
|  |  | p = 0.787 | p = 0.229 | p = 0.526 |
| Fig. 5 [A]: | *BoS<sub>w</sub>* | F<sub>(1,30)</sub> = 3.77 | F<sub>(1,94)</sub> = 0.58 | F<sub>(1,94)</sub> = 0.09 |
|  |  | p = 0.062 | p = 0.450 | p = 0.767 |
| [B]: | ln(*Percent Unstable*) | F<sub>(1,30)</sub> = 0.32 | F<sub>(1,94)</sub> = 0.00 | F<sub>(1,94)</sub> = 1.17 |
|  |  | p = 0.574 | p = 0.969 | p = 0.282 |

**Figure 4.**
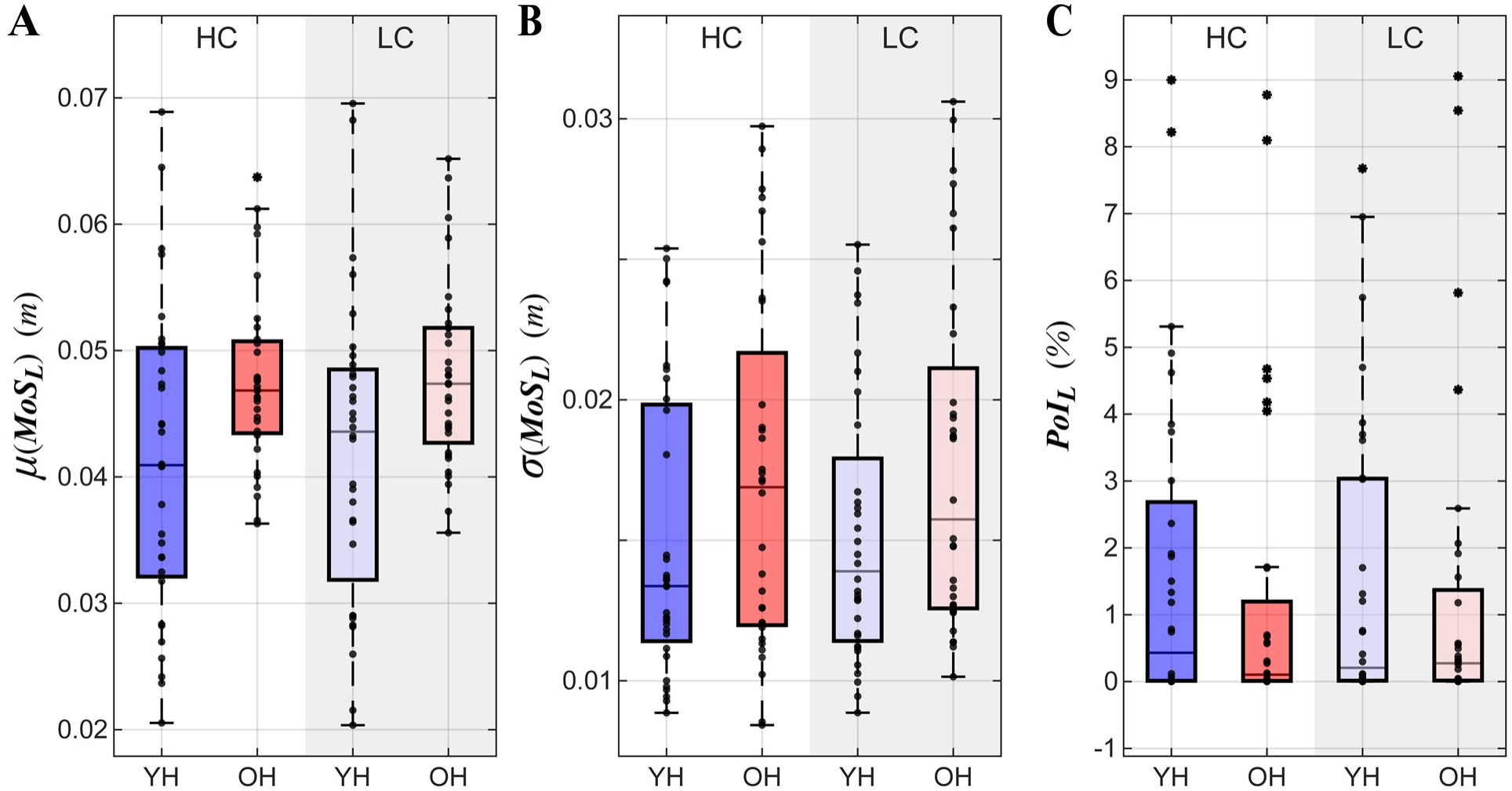
Stability Results for Walking on Straight Paths. (**A**) Lateral margin of stability means *µ*(*MoS*_*L*_), **(B)** variance *σ*(*MoS*_*L*_) and **(C)** lateral probability of instability (*PoI*_*L*_) for each Group (Young Healthy (YH); green, Old Healthy (OH); orange) and path color contrast (High (HC); light, Low (LC); dark) combination. Box plots show the medians, 1^st^ and 3^rd^ quartiles, and whiskers extending to 1.5 × interquartile range. Values beyond this range are shown as individual asterisks. The overlaid markers are individual data points for each participant from two separate trials. Detailed results of statistical comparisons are in Table 1. A significant main effect of Contrast was observed for *σ*(*MoS*_*L*_) (p = 0.008), where participants in both groups (YH and OH) exhibited greater variability when walking on the Low Contrast Straight paths.

**Table 2.** Statistical Results for Walking on Winding Paths. Differences between age Group (YH vs. OH) and path visual Contrast (HC vs. LC) for the data shown in Figs. 6 and 7.

| Data In | Variable | Group | Contrast | Group ×<br>Contrast |
|---|---|---|---|---|
| Fig. 6 [A]: | *μ*(*MoS<sub>L</sub>*) | F<sub>(1,30)</sub> = 2.75 | F<sub>(1,94)</sub> = 25.29 | F<sub>(1,94)</sub> = 1.86 |
|  |  | p = 0.108 | **p < 0.0005** | p = 0.176 |
| [B]: | ln(*σ*(*MoS<sub>L</sub>*)) | F<sub>(1,30)</sub> = 2.67 | F<sub>(1,94)</sub> = 7.91 | F<sub>(1,94)</sub> = 2.27 |
|  |  | p = 0.113 | **p = 0.006** | p = 0.136 |
| [C]: | *PoI<sub>L</sub>* | F<sub>(1,30)</sub> = 3.75 | F<sub>(1,94)</sub> = 19.55 | F<sub>(1,94)</sub> = 1.77 |
|  |  | p = 0.062 | **p < 0.0005** | p = 0.187 |
| Fig. 7 [A]: | *BoS<sub>w</sub>* | F<sub>(1,30)</sub> = 2.69 | F<sub>(1,94)</sub> = 72.38 | F<sub>(1,94)</sub> = 0.25 |
|  |  | p = 0.111 | **p < 0.0005** | p = 0.615 |
| [B]: | *Percent Unstable* | F<sub>(1,30)</sub> = 4.25 | F<sub>(1,94)</sub> = 13.09 | F<sub>(1,94)</sub> = 0.01 |
|  |  | **p = 0.048** | **p < 0.0005** | p = 0.913 |

While walking on Straight paths, older (OH) adults walked with slightly, but not significantly (p = 0.062; Table 1) wider bases of support (*BoS*_*w*_) than younger (YH) adults (Fig. 5A). Otherwise, there were no significant effects for either Group or Contrast, and no significant interaction effects for either *BoS*_*w*_ (Fig. 5A) or percent unstable steps taken (Fig. 5B).

**Figure 5.**
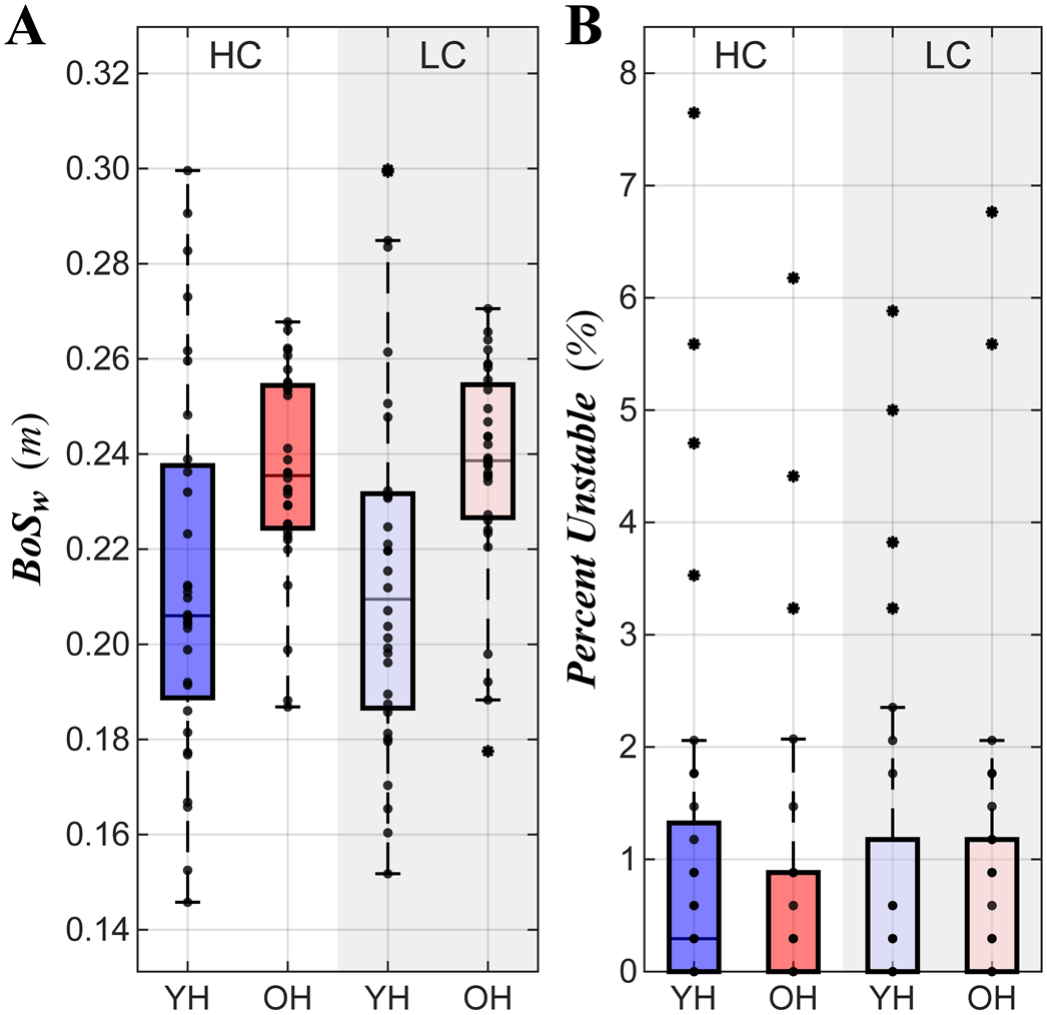
Stepping Results for Walking on Straight Paths. (**A**) Lateral base of support width (*BoS*_*w*_) and (**B**) percent of unstable steps for each Group (Young Healthy (YH); green, Old Healthy (OH); orange) and path color contrast (High (HC); light, Low (LC); dark) combination. Data are plotted in the same manner as described in Fig. 4. Detailed results of statistical comparisons are in Table 1. There were no statistically significant differences for either measure (all p > 0.06).

### Winding Paths

When walking on Winding paths, all participants in general took far more unstable (i.e., *MoS*_*L*_ < 0) steps (e.g., Fig. 3B) than they did on the Straight paths (Fig. 3A), which we expected for this more difficult task (Render et al., 2024; Render et al., 2027).

While walking on low (LC) compared to high (HC) contrast Winding paths, participants in both groups (YH and OH) exhibited smaller mean, *μ*(*MoS*_*L*_) (Fig. 6A; p < 0.0005; Table 2), but with greater variability, *σ*(*MoS*_*L*_) (Fig. 6B; p = 0.006; Table 2), which in turn produced higher *PoI*_*L*_ values (Fig. 6C; p < 0.0005; Table 2). There were no statistically significant Group differences (all p ≥ 0.062; Table 2) or Group × Contrast interaction effects (all p ≥ 0.136; Table 2) for any of these measures.

**Figure 6.**
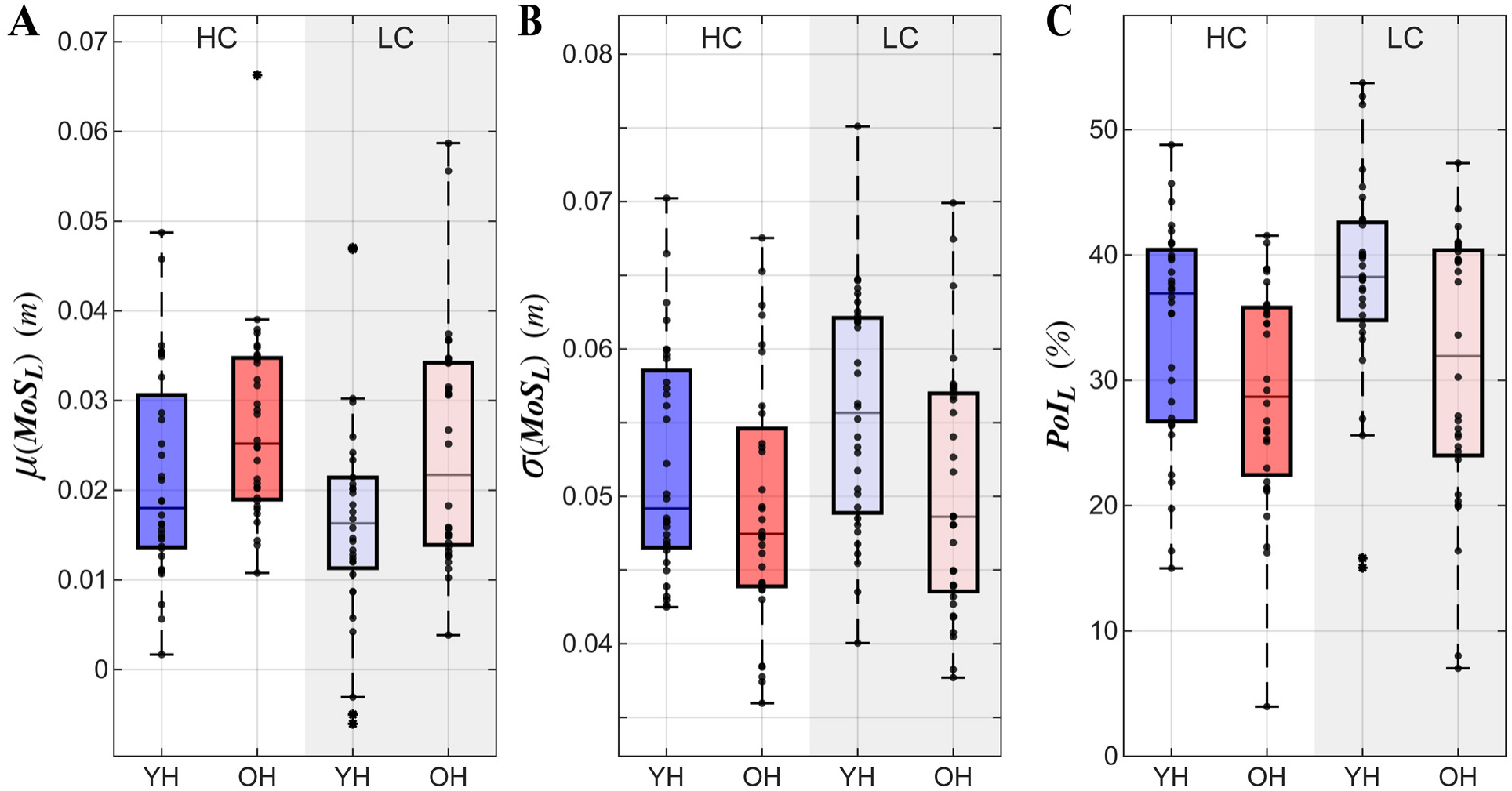
Stability Results for Walking on Winding Paths. (**A**) Lateral margin of stability means *µ*(*MoS*_*L*_), (**B**) variance *σ*(*MoS*_*L*_) and (**C**) lateral probability of instability (*PoI*_*L*_) for each Group (young healthy (YH); green, old healthy (OH); orange) and path color contrast (High (HC); light, Low (LC); dark) combination. Data are plotted in the same manner as described in Fig. 4. Detailed results of statistical comparisons are in Table 2. Significant main effects for Contrast were observed for all variables (p < 0.0005), with participants in both groups (YH and OH) exhibiting smaller *µ*(*MoS*_*L*_), but greater *σ*(*MoS*_*L*_) and larger *PoI*_*L*_ when walking on the Low Contrast Winding paths.

Likewise, while walking on low (LC) compared to high (HC) contrast Winding paths, participants in both groups (YH and OH) walked with narrower steps, (*BoS*_*W*_) (Fig. 7A; p < 0.0005; Table 2), and took significantly more unstable steps (Fig. 7B; p < 0.0005; Table 2). In addition, older (OH) adults took significantly fewer unstable steps than young (YH) adults (Fig. 7B; p = 0.048; Table 2). There were no significant Group × Contrast interaction effects (all p ≥ 0.615; Table 2) for either of these measures.

**Figure 7.**
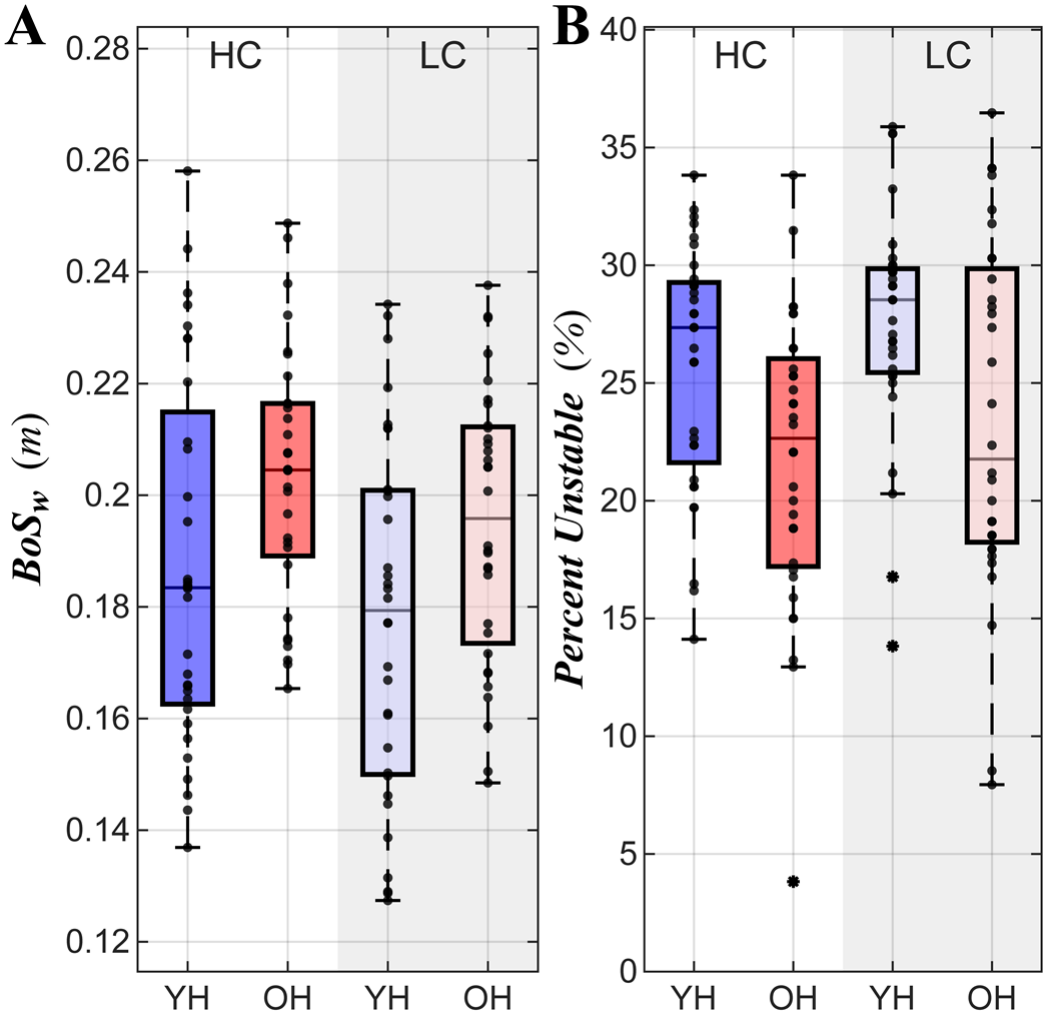
Stepping Results for Walking on Winding Paths. **A**) Lateral base of support width (*BoS*_*w*_) and **(B)** percent of unstable steps for each Group (Young Healthy (YH); green, Old Healthy (OH); orange) and path color contrast (High (HC); light, Low (LC); dark) combination. Data are plotted in the same manner as described in Fig. 4. Detailed results of statistical comparisons are in Table 2. Significant main effects for Contrast were observed for both variables (p < 0.0005), with participants in both groups (YH and OH) taking steps with narrower *BoS*_*w*_, but larger *PoI*_*L*_ when walking on the Low Contrast Winding paths. A significant Group effect was also found for *BoS*_*w*_ (p = 0.048), with OH participants walking with slightly wider *BoS*_*w*_ than YH participants.

## DISCUSSION

To avoid falling in everyday life, individuals must continuously maintain their balance as they navigate complex environments. Gait is shaped not only by neuromuscular control but also by visual input (Muller et al., 2024; Tong, 1995), which helps guide foot placement and avoid obstacles. With aging, declines in visual acuity and sensory integration can alter both perception and movement strategies. This study quantified how aging affects mediolateral stability when walking on real-world-like paths of varying complexity and visual contrast.

On Low (LC) contrast Winding paths, participants in both groups (YH and OH) took narrower steps (*BoS*_*w*_; Fig. 7A), contributing to decreased *μ*(*MoS*_*L*_) (Fig. 6A). It’s possible they did so to reduce visual uncertainty so they could focus on task-relevant visual information (Domínguez-Zamora and Marigold, 2021; Tong et al., 2017). However, the reduced visibility of these walking paths also led people to take more variable steps (*σ*(*MoS*_*L*_); Fig. 6B), which, in combination with their decreased *μ*(*MoS*_*L*_), contributed to greatly increased *PoI*_*L*_ (Fig. 6C) and taking more unstable steps (Fig. 7B). Taken together, these findings demonstrate that on these harder-to-see LC Winding paths, participants chose to take narrower steps to avoid stepping off their path, despite such narrower steps increasing their risk of lateral instability (Kazanski et al., 2024).

These effects were far less pronounced when participants walked on Straight paths (Figs. 4-5). YH and OH participants both exhibited greater step-to-step variability in *MoS*_*L*_ on Low (LC) compared to High (HC) paths (Fig. 4B), but otherwise exhibited no significant differences between Groups or Contrasts (Table 2). Hence, walking on non-straight paths is both biomechanically more demanding (Render et al., 2024, 2025) and *visually* more demanding (Matthis et al., 2018; Render et al., 2027; Tong et al., 2017).

As we observed previously (Render et al., 2024), the step-to-step distributions of *MoS*_*L*_ appeared somewhat skewed in the Winding path trials (e.g., Fig. 3B). We therefore correlated our *PoI*_*L*_ estimates (Figs. 4C and 6C) with the directly-calculated % Unstable Steps (Figs. 5B and 7B). As anticipated (Render et al., 2024), *PoI*_*L*_ estimates somewhat over-estimated the actual % Unstable steps taken, particularly on the Winding paths (Fig. 8A). Nevertheless, both measures were highly correlated (*r*^2^ > 80%), regardless of path. This further substantiates that *PoI*_*L*_ is a good proxy for quantifying the likelihood of taking unstable steps in contexts where, for example, it may not be feasible to record large numbers of steps (Render et al., 2024).

**Figure 8.**
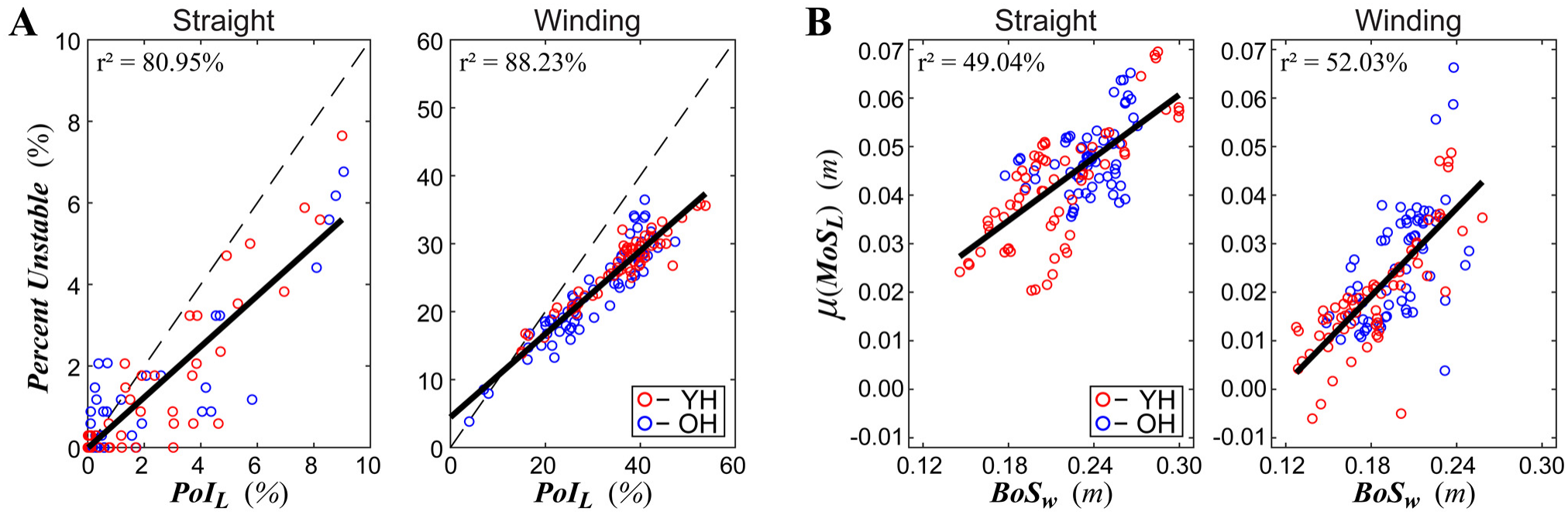
Correlations. (**A**) Lateral probability of instability (*PoI*_*L*_) slightly under-estimated, but was strongly correlated (r^2^ > 80%) with Percent Unstable Steps, when participants walked on either Straight or Winding paths (note the scales on both sets of axes are different). (**B**) Base-of-support width (*BoS*_*w*_) was strongly correlated (r^2^ > 49%) with average margin of stability (<u>μ</u>(*MoS*_*L*_)), whether participants walked on either Straight or Winding paths (note the scales on both sets of axes are the same). For all plots, data were pooled across all trials and all visual conditions. Individual markers represent individual trials for both young (YH; red) and older (OH; blue) participants.

It has been widely observed that average measures of *MoS*_*L*_ are larger (i.e., seemingly *more* “stable”) when healthy adults are subjected to *de*-stabilizing perturbations (McAndrew Young and Dingwell, 2012; Onushko et al., 2019), and in fall-prone populations (Watson et al., 2021). To assess this, we directly correlated *BoS*_*w*_ with *μ*(*MoS*_*L*_), and indeed, *BoS*_*w*_ was a very strong contributor (*r*^2^ > 49%) to *μ*(*MoS*_*L*_) across groups and conditions (Fig. 8B). Participants who adopted wider bases of support (*BoS*_*w*_) may have been adopting a more conservative gait strategy intended to mitigate their risk of losing balance (Kazanski et al., 2024).

Contrary to our hypothesis however, we observed few differences between the young (YH) and older (OH) adults tested here. Our OH adults exhibited slightly (but significantly) elevated TUG and FSST scores (Table 1). However, these remained well below values typical of impaired mobility for both TUG (Bohannon, 2006; Gatenio-Hefling et al., 2025) and FSST (de Aquino et al., 2022; Whitney et al., 2007). Likewise, our OH adults exhibited similar self-selected walking speeds in the 5mWT (Table 1), further indicating that our OH adults were substantially high-functioning (Middleton et al., 2015).

Indeed, while these OH adults tended to take slightly wider steps (*BoS*_*w*_; Figs. 4A and 6A), these differences did not reach statistical significance (Tables 1-2). Moreover, our OH adults exhibited *lower* % Unstable Steps (Fig. 7B), resulting in lower *PoI*_*L*_ (Fig. 6C) than YH adults when walking on Winding paths. Taken together, these findings suggest that healthy older adults can successfully adapt their gait to maintain lateral stability, even when walking conditions become more visually and mechanically challenging. Future studies, however, are needed to determine if the present findings can generalize to populations less able to adapt to such walking challenges, particularly to more frail and/or visually impaired older adults.

A central question this study addressed was whether lower visual contrast disproportionately impacts lateral stability in older adults. Despite well-documented age-related declines in visual processing (Madden, 2007; Owsley, 2011), no Group × Contrast interactions were observed for any of the primary stability metrics. Thus, the OH adults tested here were not more adversely affected by low-contrast walking conditions than our YH adults. However, complementary analyses on this dataset (Overby et al., 2027) did reveal significant Group × Contrast interactions in other stepping regulation metrics.

Those findings suggest that perhaps these OH adults’ stable outcomes reported here resulted from them adopting effective compensatory adjustments in stepping. However, future work should test this more explicitly.

This study offers important implications for how aging affects balance control during walking. First, our findings suggest that *PoI*_*L*_ quantifies dynamic balance risk better than *μ*(*MoS*_*L*_) by capturing how stability fluctuates across steps. Second, our findings suggest specific ways interventions aimed at fall prevention might benefit by enhancing adaptive stepping strategies that exploit perceptual-motor integration. In particular, directly manipulating visual uncertainty about stepping/walking task goals challenges participants to focus on *task*-relevant visual information (Domínguez-Zamora and Marigold, 2021; Tong et al., 2017). This would likely encourage balance-impaired older adults to strengthen their visual-motor coordination to reinforce compensatory visual sampling (Dowiasch et al., 2015; Meissner et al., 2025) and stepping behaviors to reduce their risk of losing balance.

## ACKNOWLEDGEMENTS

The authors thank Dr. Anna C. Render and Ethan Friedman for their contributions and technical support throughout data collections. This work was partly supported by NIH grant 1-R01-AG049735 (to JBD).

